# Stability of Ecologically Scaffolded Traits During Evolutionary Transitions in Individuality

**DOI:** 10.1101/2023.08.17.553478

**Authors:** Guilhem Doulcier, Peter Takacs, Katrin Hammerschmidt, Pierrick Bourrat

## Abstract

Evolutionary transitions in individuality (ETIs), such as the emergence of multicellularity, are events in the history of life during which entities at one level of organisation (particles) form collective-level entities that subsequently become individuals in their own right. Recent empirical and theoretical studies advocate the importance of an externally imposed meta-population structure or “ecological scaffold” for the emergence of new levels of individuality. Such a scaffold enables survival and reproduction at the collective level and thus the possibility of selection for beneficial traits on that level. However, a long-standing difficulty for the ecological scaffolding approach has been its inability to adequately explain how collective-level trait values that evolved under scaffolding conditions can be retained once these conditions are lifted. We call this difficulty “the problem of endogenisation.” Here, we derive general conditions for the possibility of endogenisation. Key to endogenisation is the existence of a fitness valley that can be circumvented when scaffolding occurs. Using a stochastic meta-population model, we implement two versions of ecological scaffolding (one temporal and one spatial) and study subsequent evolutionary trajectories using the modelling techniques of adaptive dynamics. Our analysis yields several important results. The temporal model reveals that only collective traits based on particle-particle interactions can be endogenised when a temporary scaffold is applied to the entire population. The spatial model shows that, given the presence of an environmental gradient of externally imposed meta-population structure, ecological scaffolding can only occur in a limited “Goldilocks” zone of the environment. Further, if endogenisation conditions are also fulfilled, scaffolded collectives can colonise non-scaffolding areas of the environment. We conjecture that Goldilocks zones could act as initiators of ETIs and help explain the near ubiquity of collective-level individuality even if the conditions that promote it prove to be rare.

## Introduction

Evolutionary transitions in individuality (ETIs) involve the emergence of collective-level individuals from independently reproducing particle-level individuals [1–11]. The evolution of multicellularity is considered a prime example of an ETI [9, 12, 13]. The emergence of multicellular organisms [14], organelles through endosymbiosis [15] and eusocial organisations [16] have occurred several times in the history of life. Multiple independent occurrences of similar transitions strongly suggest that there may be general mechanisms promoting ETIs.

Recently, the “ecological scaffolding” scenario has been proposed as a possible mechanism for ETIs [17–19]. The ecological scaffolding scenario is one among several mechanisms for ETIs that have been proposed in the literature [2, 4, 20, 21]. Key to this model is that an ETI can be initiated from a specific kind of externally imposed meta-population structure—a *scaffold*—consisting of a set of populations of particles confined to bounded niches with limited dispersal, ranging from physical structures to organisms hosting symbionts (Figure 1, arrow 1). In situations where niches can be depleted and regenerated (newly available structures, surfaces, or hosts), the colonisation of new niches and the extinction of existing populations creates a birth-death process at the collective (population) level. Under specific conditions, this birth-death process provides an opportunity for natural selection to promote beneficial traits at the collective level (Figure 1, arrow 2).

**Fig. 1.**
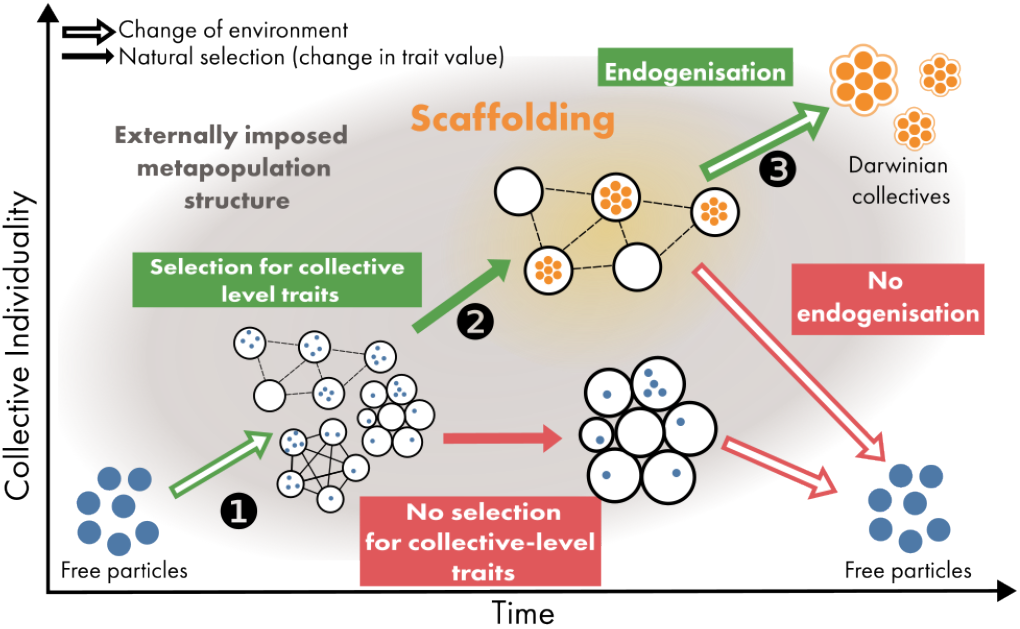
Endogenisation of scaffolded traits. Scaffolded traits evolve when the population is structured in a specific way. When the meta-population structure is modified (i.e., when the scaffold is removed), the properties can either revert to their ancestral values (no endogenisation) or keep the same values (endogenisation). Under the ecological scaffolding model, an ETI involves the formation of an externally imposed meta-population structure from free particles (arrow 1). Trait *scaffolding* is the selection of a trait value in specific conditions of the environment (i.e., scaffolding environment, arrow 2) that promotes survival and reproduction at the collective level. A scaffolded trait is said to be *endogenised* if it is stable even when the meta-population conditions that were required for its emergence are perturbed (arrow 3).

Ecological scaffolding has been shown to successfully select for traits associated with higher collective-level persistence and fertility, such as: lower particle growth rate in a limited resource environment [17], particle-particle interaction that ensures transgenerational stability of the collective composition [22], and proto-germ-soma differentiation [11].

Although they provide a simple mechanism for the emergence of collectives and traits at that level, previous articulations of the ecological scaffolding model are nevertheless insufficient. In nature, collective-level entities that are considered individuals typically do not exhibit an external scaffold. Whatever scaffolds there may once have been, collective-level individuals have done away with them over time, just as a successfully constructed building can stand true after the scaffolding that supported it has been removed. Thus, it appears that an ETI is more fully realised when the selected collective-level traits persist even when the scaffold is lifted (Figure 1, arrow 3). This phenomenon is referred to as *endogenisation* [17, 19].

Scaffold endogenisation remains an all-important but un-solved issue for this theory. Explanations to date have been unable to provide convincing reasons for differentiating cases where collective-level entities persist as individuals from cases where they would disintegrate were it not for the presence of scaffolding conditions. The standing challenge is to provide an adequate mechanistic explanation of endogenisation. We address this challenge by delineating the conditions under which a scaffolded collective-level trait can remain evolutionarily stable even when the scaffold is lifted. The idea of “lifting the scaffold” can be implemented in several ways. While it could be implemented by removing all externally imposed meta-population structures (as in [17, 19]), we here assume that an externally imposed meta-population structure is ever-present and instead choose to adjust its defining parameters so that scaffolded traits would not have evolved in the first place (i.e., by reverting them to pre-scaffolding parameter values).

To do so, we propose a general mechanistic model (outlined in Sections A and B) for a wide range of meta-population structures and derive formal conditions (Section C) under which ecological scaffolding can lead to endogenisation once the scaffold is lifted. Using this model, we show how a collective-level trait can be endogenised (Section D). We then explore some consequences of this phenomenon for ETIs. One particularly noteworthy consequence is that the specific environmental conditions under which collective individuals emerge does not fully determine their capacity to colonise new niches (Section E).

## Results

### A. Stochastic meta-population model of ecological scaffolding

To explore a wide range of scaffolding situations, we model a meta-population of a fixed number *D* of niches connected on a graph, each initially containing *R* re-sources, and particles that each carry a mutable trait *θ*. Particles consume resources at a rate 1 and, upon consuming resources, either duplicate or become a propagule with probability *p* and (1 − *p*), respectively (Figure 2a). When a niche is out of resources, all particles within it die instantly and resources are replenished to the initial value *R*. The probability *p* depends on the trait *θ* and on the local state of the niche (predominantly, the number of particles within it). We study two functional forms for the dependency of *p* on *θ*, which are detailed in Sections B and D. When a particle migrates out of a niche, it colonises an empty connected niche. If no such niche exists, the propagule-particle dies out. These rules constitute a Markov jump process with events detailed more formally in Supplementary Section 3. As this stochastic process is a continuous-time Markov jump process, it can be approximated by a set of ordinary differential equations (ODE), as detailed in Supplementary Note 1. The simplified ODE system has two key parameters for a given trait *θ* value, the life-time of a collective *τ* (*θ*) and the lifetime expected number of propagules for a collective *ρ*(*θ*). These two values can be derived from the stochastic system, as shown in Supplementary Note 2. The analysis of the simplified ODE system and the derivation of the fitness invasion gradient is presented in Supplementary Note 1.

**Fig. 2.**
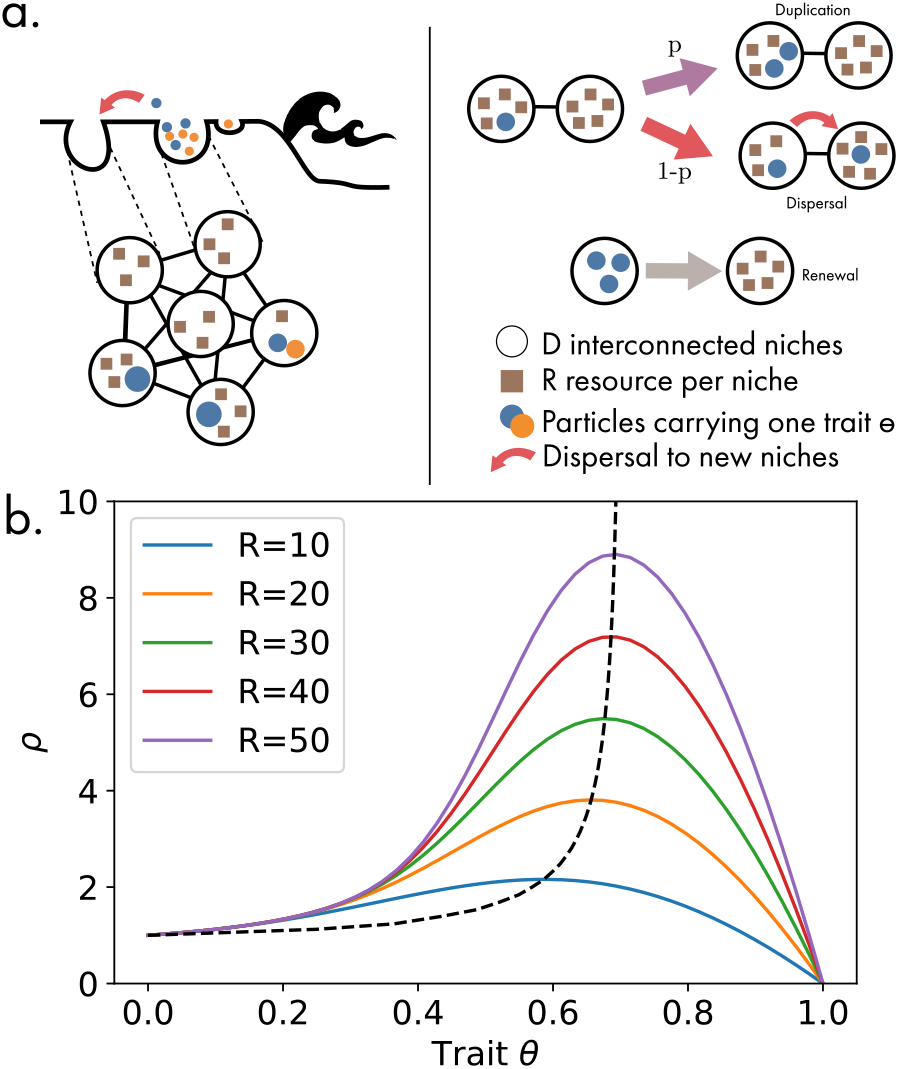
Evolutionary dynamics. **(a) Meta population model**: In our model, there are D available niches that initially contain *R* resources. Particles reproduce or migrate with rate 1. Particles carry one trait *θ* that determines the probability that the particle will reproduce (*p*(*θ*)) or migrate to an empty connected niche (1 − *p*(*θ*)). When all resources are consumed, all particles within the niche die and the re-sources are replenished to the initial value R. **(b) Intrinsic duplication-dispersal ratio**: When the probability of dispersal or duplication is intrinsic and constant (*p*(*θ*) = *θ*), there is a global optimal trait value for each resource level. If the meta-population changes, the trait value tracks the new optimum, but there is no endogenisation.

Using this general framework, we study the problem of endogenisation. Following previous work [17, 19], endogenisation of a trait occurs when the scaffolded value persists after the scaffolding conditions revert to pre-scaffolding conditions. First, we demonstrate that a simple trait value (i.e., taking a constant intrinsic value for the duplication-to-dispersal value, *p* = *θ*) can change as the scaffolding meta-population reverts to a non-scaffolding state (Section B). This shows that not all scaffolded traits are necessarily endogenised (i.e., stable when the environment is reverted to pre-scaffolding conditions). Thus, we establish formal conditions for the endogenisation of traits based on the structure of the trait-fitness landscape (Section C). We then choose a trait fulfilling these formal conditions (i.e., taking a duplication-to-dispersal ratio that is sensitive to local population density) and show that its value is stable even if the scaffolding conditions are lifted (Section D). Finally, given a gradient in externally imposed meta-population structure, we show how collective-level entities can colonise new areas through endogenisation despite the restriction of scaffolding to limited zones of the environment. These zones can effectively become initiators of ETIs through a phenomenon we call the “Goldilocks Zone Effect” (Section E).

### B. An intrinsic particle dispersal-duplication ratio evolves in response to the scaffold but is not endogenised

Here, we examine the simplest functional form for the trait and study its evolutionary response to changes in the meta-population parameter. Consider a case where the mutable trait *θ* is carried by each cell and directly encodes the rate at which cells duplicate (*p*) or leave the niche (1 − *p*), such that *p*(*θ*) = *θ*, independently of their surroundings. This trait encodes the investment towards particle duplication or dispersal in the shape of a linear tradeoff. At the collective level, this translates into a non-linear tradeoff between the life span of collectives *τ* (*θ*) and the average number of propagules they produce *ρ*(*θ*) (due to demographic effects, as illustrated in Supplementary Figure 5). Indeed, the limited resources in the niches are allocated either to producing more particles or to producing propagules. Let us detail the behaviour of collectives for different values of *θ*.

Note that the two extreme cases, *θ* = 0 and *θ* = 1, are not viable. First, consider the ecological dynamics of a unique niche seeded with one cell. If *θ* = 0, particles never reproduce and always move from one niche to the next. Thus, in this limited case, the lifetime reproductive output of a niche is always *ρ*(0) = 1 and its life span is, on average, *τ* (0) = 1. The populations of particles and of collectives never grow in size because no particle reproduction occurs. Conversely, particles never become propagules if *θ* = 1. They duplicate until all resources are expended and the local population goes extinct. Thus, *ρ*(1) = 0. Intermediate values of *θ* have higher values of *ρ*, as shown in Figure 2b.

Now, if we consider the ecological dynamics of the entire meta-population, we know from the analysis of the ODE system that the population is only viable if *ρ*(*θ*) *>* 1 and reaches an equilibrium value that depends on the number of resources in a niche *R* (see Supplementary Figure 1 for the equilibrium values as a function of *R*). Thus, the population is viable for values of *θ* strictly higher than zero and smaller than a threshold 0 *< θ*_max_ *<* 1, such that *ρ*(*θ*_max_) = 1.

To predict the evolutionary trajectories of the system, we turn to invasion analysis. Consider a homogeneous population at ecological equilibrium. The fate of a single new mutant particle with a different value of *θ* in a new niche can be predicted using the ODE model. It has either a positive growth rate (black zone in the pairwise invasibility plot, Supplementary Figure 2), and thus a possibility to become fixed in the meta-population, or a negative growth rate (white zone, Supplementary Figure 2). The value of the invasion fitness results in an evolutionarily stable strategy (ESS; *θ*^∗^) that is convergence-stable (i.e., a resident population with a trait value that is close but not exactly equal to the ESS can be invaded by mutants with an even closer trait value to the ESS) and evolutionarily stable (i.e., a population with the trait value *θ*^∗^ cannot be invaded by mutants with a close trait value). The value of the ESS depends on the number of re-sources *R* in a single niche.

Finally, consider the case in which *θ* can mutate freely, starting from an ancestral value *θ*_0_ that is viable (0 *< θ*_0_ *< θ*_max_). When mutations are so rare that populations reach a new ecological equilibrium between the emergence of new mutants (i.e., the ecological and evolutionary time scales are separated), adaptive dynamics predict that the meta-population will converge towards the evolutionary stable strategy. An example of a stochastic trajectory is shown in Supplementary Figure 6. The value of the ESS for each value of *R* is shown as a dashed line in Figure 2b and Supplementary Figure 3.

In this model, the selected trait value depends on the meta-population parameters (here, the number of resources in a niche). If the parameter *R* changes, the selected trait value simply tracks the optimal probability associated with the change, which means there is no endogenisation.

### C. Formal conditions for trait endogenisation: circumvention of a fitness valley

Having shown that scaffolding does not entail endogenisation, we now establish the conditions for endogenisation. A formal derivation of these conditions can be found in Supplementary Section 5.

Consider a two-dimensional landscape that maps the state of the environment and the value of a collective trait to its fitness in this environment. This simplified landscape (illustrated in Figure 3) allows us to describe the conditions in which scaffolding and endogenisation are possible. To keep the model simple, we consider only two states of the environment— namely, “scaffolding” and “non-scaffolding”—as well as two collective trait values—namely, “no collective organisation” and “collective organisation.” This results in four trait-environment combinations. By “collective organisation,” we mean that the trait exhibits a behaviour different from the intrinsic duplication-dispersal trait, or what we refer to as “non-aggregativity” (see Discussion).

**Fig. 3.**
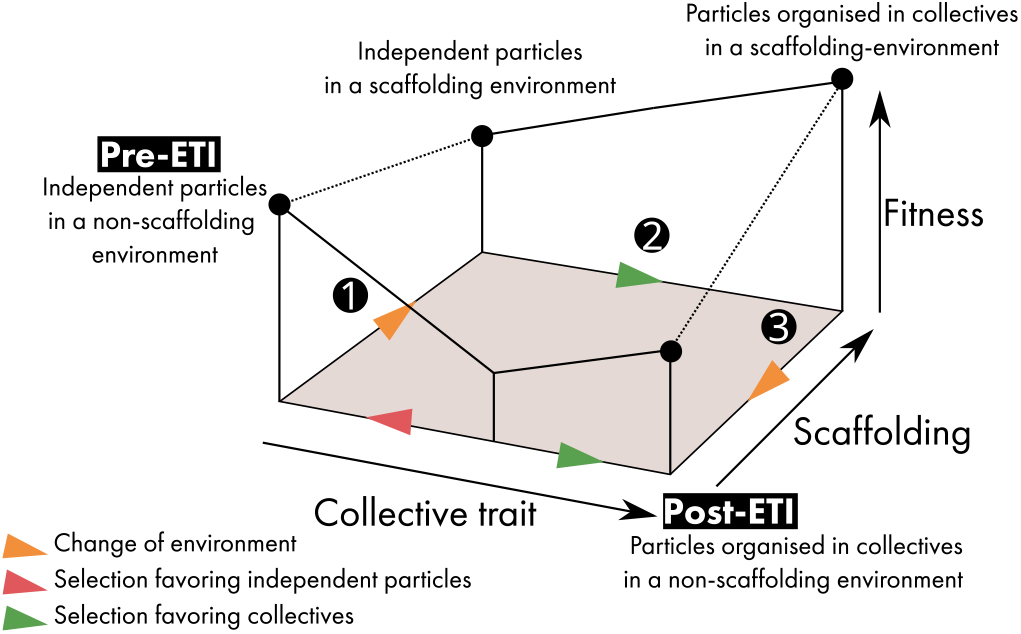
Conditions for endogenisation. A simplified scaffolding scenario in three steps: **1**. Change in the environment **2**. Selection of the collective trait in the scaffolded environment (Scaffolding) **3**. Change in the environment. A collective trait is said to be *scaffolded* if a change in the population structure results in its evolution by natural selection (Steps 1–2). A collective trait is said to be *endogenised* if it is evolutionarily stable after the scaffolding conditions are lifted (Step 3).

In the initial condition, we assume that the environment is non-scaffolding and there is no collective organisation. This state is evolutionarily stable. A scaffolding experiment is then conducted. At time 0, the environment is changed to a scaffolding condition (Step 1), held constant for a duration *T* (Step 2), and then returned to the non-scaffolding condition (Step 3).

A collective trait is said to be *scaffolded* if a change in the population structure results in the evolution by natural selection of a collective-level trait (Steps 1–2). A collective trait is said to be *endogenised* if it is evolutionarily stable after the scaffolding conditions are lifted (Step 3).

This scaffolding-endogenisation trajectory (Steps 1–3) is only possible if three conditions are fulfilled. First, both trait values must be ESS in the non-scaffolding environment. Second, the “no collective organisation” trait value must be evolutionarily unstable in the scaffolding environment. Third, the trait value of the population after a duration *T* must fall within the basin of attraction for the trait “collective organisation” in a non-scaffolding environment. In other words, the transition to a scaffolding environment from a non-scaffolding environment and then back again allows populations to circumvent a fitness valley that exists only in the non-scaffolding environment and would prevent evolution towards collective organisation if no scaffold was imposed.

These conditions are not fulfilled in the simple model without the particle-particle interactions presented in Section B. Indeed, there is a single ESS for each value of *R*. There is consequently no fitness valley in the non-scaffolding environment to be circumvented. Thus, some interactions between particles are necessary for endogenisation to be successful in this model.

In the next two sections, we follow a trait that fulfils the endogenisation conditions. In the first section, we study the evolutionary stability of the derived trait value in the face of environmental change. In the second section, we explore the consequences of this stability when the population is put in an environmental gradient.

### D. A density-dependent particle dispersal-duplication ratio fulfils the conditions for endogenisation

#### D.1. The optimal phenotype for resource use is to keep the population size as small as possible

In this section, we present a variation of the model in which the probability for a particle to either duplicate or disperse depends on its ability to sense the presence of other particles in the niche. This model with particle-particle interactions fulfils the endogenisation conditions outlined in Section C. There is a global maximum of fitness that corresponds to a particle that always duplicates if it is alone and always disperses if that is not the case. This scenario results in the maximum possible value of *ρ* = *R/*2. This phenotype is a fitness maximum due to two simplifying assumptions of the system.

First, we assume that there is no particle death outside of starvation events (when all the resources in a niche are depleted). As a consequence, the most efficient proto-organisms can get away with a “soma size” of one particle (because a bigger investment in soma does not translate into an increase in viability) and invest the rest of the resources in fertility. Introducing density-dependent collective death (modelling predation or other external perturbations) would relax this assumption, and the optimal soma size would likely be higher than one particle. However, this would complexify the model without bringing more clarity to the concept of endogenisation.

Second, our model assumes that propagules can only colonise empty niches. There is no advantage in a large particle population within the niche. Although several phenotypes can compete within a single niche, there is a bottleneck of one particle when niches are newly colonised.

#### D.2. Model for the evolution of a sensory system

Consider a particle with the optimal strategy (always duplicate if alone in the niche and always disperse otherwise) but with a flawed capacity to assess its surroundings, particularly the number of other particles within the niche. The emergence of a sensory (chemical or physical) system might be an instance of this insofar as it could initially bias the probability of activating the molecular pathways leading to duplication or dispersal. To model such a case, let the mutable trait *θ* be the “error of the sensory system” (see Figure 4a), *n* be the true number of particles within the niche, and let the probability for a particle to duplicate be:

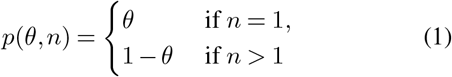

**Fig. 4.**
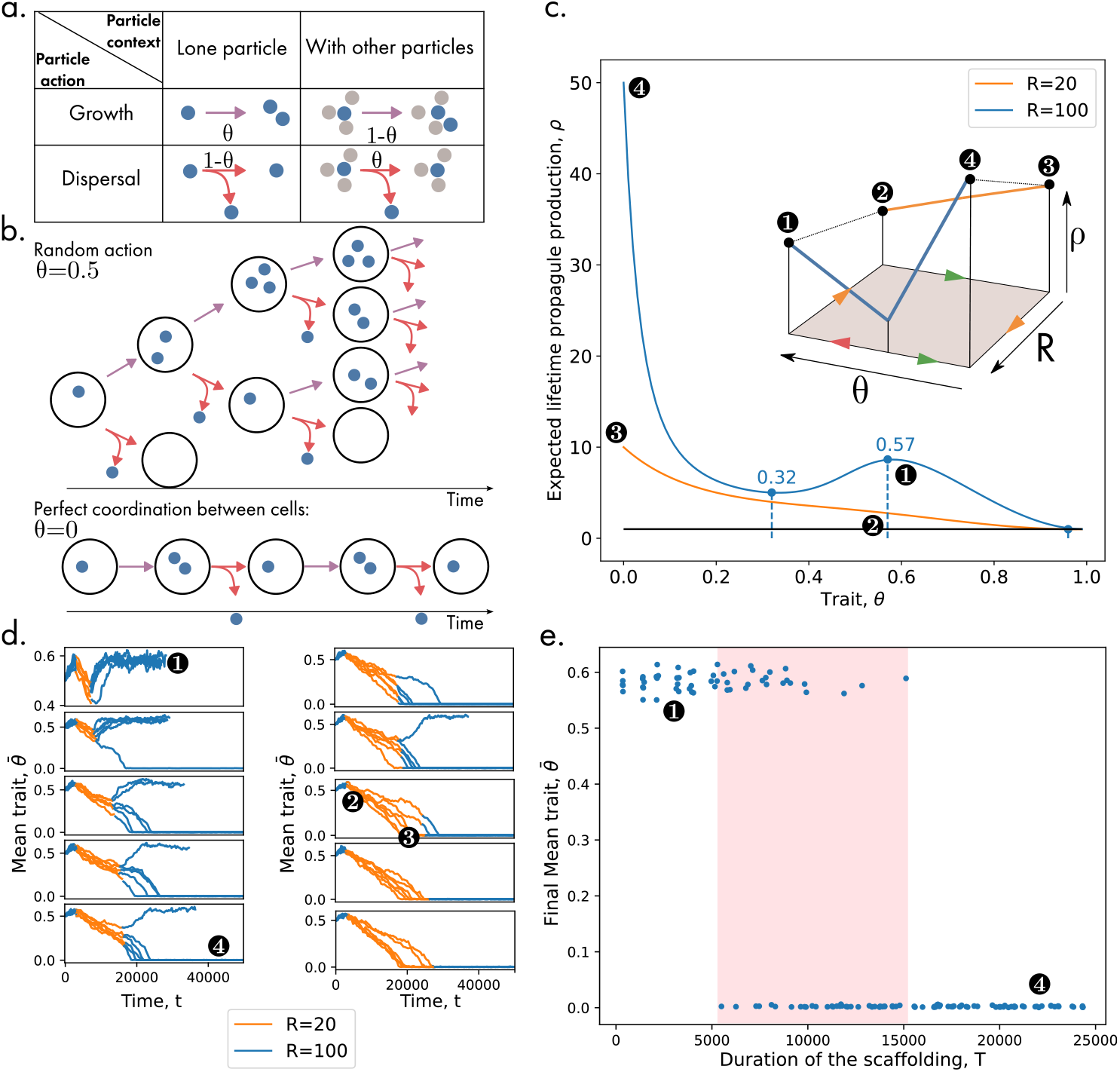
Endogenisation of a scaffolded trait value. **a**. *Model*. In the density-dependant duplication-dispersal ratio, the trait value *θ* controls the probability to duplicate rather than disperse when the particle is alone and, conversely, to disperse rather than duplicate when the particle is not alone in the niche. If *θ* = 0.5 (ancestral trait value), particles act the same way regardless of the local density of particles as in the intrinsic ratio model (Figure 2) with value 0.5. **b**. *Extreme traits values*. The optimal value of *θ* is 0 as it produces the most propagules without wasting any resources (e.g., by leaving a niche before all resources are expended). **c**. *Propagule production as a function of the trait value*. In larger niches (*R >* 39), there is a fitness valley between the ancestral value 0.5 and the coordinated value 0. This valley does not exist in small niches. Hence, the trait value fulfils the conditions for endogenisation. **d**. *Evolutionary trajectories*. If particles from large niches are put into smaller niches for a limited duration (orange part of the trajectory), the average trait value tends towards 0 as lower values are selected. However, if they are put back in larger niches too soon, they revert to the ancestral value. **e**. *Evolutionary endpoint as a function of the duration of scaffolding by small niches*. Intermediary scaffolding durations (red area) do not always lead to endogenisation because of stochastic effects.

If *θ* = 0, the sensory system is perfect, and the particle always duplicates if it is alone (*p*(0, 1) = 1) and disperses otherwise (*n >* 1, *p*(0, *n*) = 0) (see Figure 4b). If *θ* = 0.5, the sensory system is non-functional, and the probability for a particle to duplicate or disperse is the same regardless of the true state of the niche (see Figure 4b). In the following, we assume that the ancestral state is that cells have no sensory system, *θ*_0_ = 0.5 (which makes the phenotype equivalent to the one in Figure 2 with *θ* = 0.5), and ask whether better sensory systems (lower values of *θ*) can evolve due to scaffolding and be endogenised in the sense of Section C. If *θ* = 1, the sensory system is perfectly accurate but translates to a non-viable strategy where particles always disperse to new niches without ever duplicating. Finally, values of *θ* between 0 and 0.5 (and between 0.5 and 1) correspond to different degrees of accuracy of the sensory system.

As before, we again consider just two environments: non-scaffolding and scaffolding. However, the environment is here deemed to be non-scaffolding when it is composed of large niches (*R* = 100), whereas an environment composed of smaller niches (*R* = 20) is regarded as a scaffolding one. As shown in Figure 4c, this setup meets the three conditions spelled out in Section C. It presents an imperfect sensory system (*θ*_*u*_ = 0.57) that is an ESS in the non-scaffolding environment and a perfect sensory system (*θ*_*s*_ = 0), the latter of which is both (i) the only ESS in the scaffolding environment and (ii) an ESS in the non-scaffolding environment. The ESS associated with the perfect sensory system is also clearly separated from the other ESS by a valley of fitness (with a minimum in *θ* = 0.32).

#### D.3. The duration of the scaffolding must be sufficiently long to allow the crossing of the fitness valley

Given the foregoing setup, we begin by allowing an initial population of particles with ancestral trait *θ*_0_ to evolve in the non-scaffolding environment for a duration *T*_0_. The value of the environment is then changed so that it satisfies the conditions for scaffolding for a duration *T*. Finally, we revert the environment back to the non-scaffolding value.

Figure 4d shows simulations of such temporary scaffolding experiments. Initially (before *T*_0_), the populations evolve towards a mean trait value of *θ*_*u*_ *> θ*_0_. When the scaffolding is applied, the evolutionary trajectory shifts towards *θ*_*s*_ = 0. If the scaffolding is lifted after *θ*_*s*_ has been reached, the mean trait in the population is stable and endogenisation has occurred. However, if the scaffolding is lifted before *θ*_*s*_ has been reached, the population mean trait could revert back to *θ*_*u*_, depending on whether there were enough individuals in the population that had a value of *θ* low enough to have crossed the fitness valley between *θ*_*u*_ and *θ*_*s*_.

Figure 4e shows the proportion of “successful” scaffolding for different values of *T*. Note that intermediate values of *T* do not always result in endogenisation. This non-deterministic behaviour comes from the stochastic nature of the mutations that do not occur exactly at the same moment in replications of the simulation.

Overall, the density-dependent particle dispersal-duplication ratio studied in this section fulfils the three conditions for endogenisation and, accordingly, displays a corresponding evolutionary trajectory: when niche size is reduced, coordination between particles is selected. Coordination is retained (i.e., is evolutionarily stable) even when niche sizes return to their ancestral values.

### E. Meta-population structure gradients can feature “Goldilocks zones” as ETI initiators

In this section, we replace the temporary scaffolding (changing the environment for a limited duration) with a standing environmental gradient that features both scaffolding and non-scaffolding conditions. Consider a diversity of niches along a size gradient from very large niches (*R* = 100) to very small niches (*R* = 2, see Figure 5a). There are 10 niches for each resource richness class and 98 different classes, for a total of *D* = 980 niches. Dispersal is only allowed between niches of the same or adjacent richness (e.g., particles in a niche *R* = 50 can disperse towards niches with *R* = 50, *R* = 49 or *R* = 51). Initially, ancestral particles with *θ*_0_ = 0.5 are seeded in the richest niches (*R* = 100) and the dynamic of the population is simulated. Figure 5b–d shows the result of such a simulation.

**Fig. 5.**
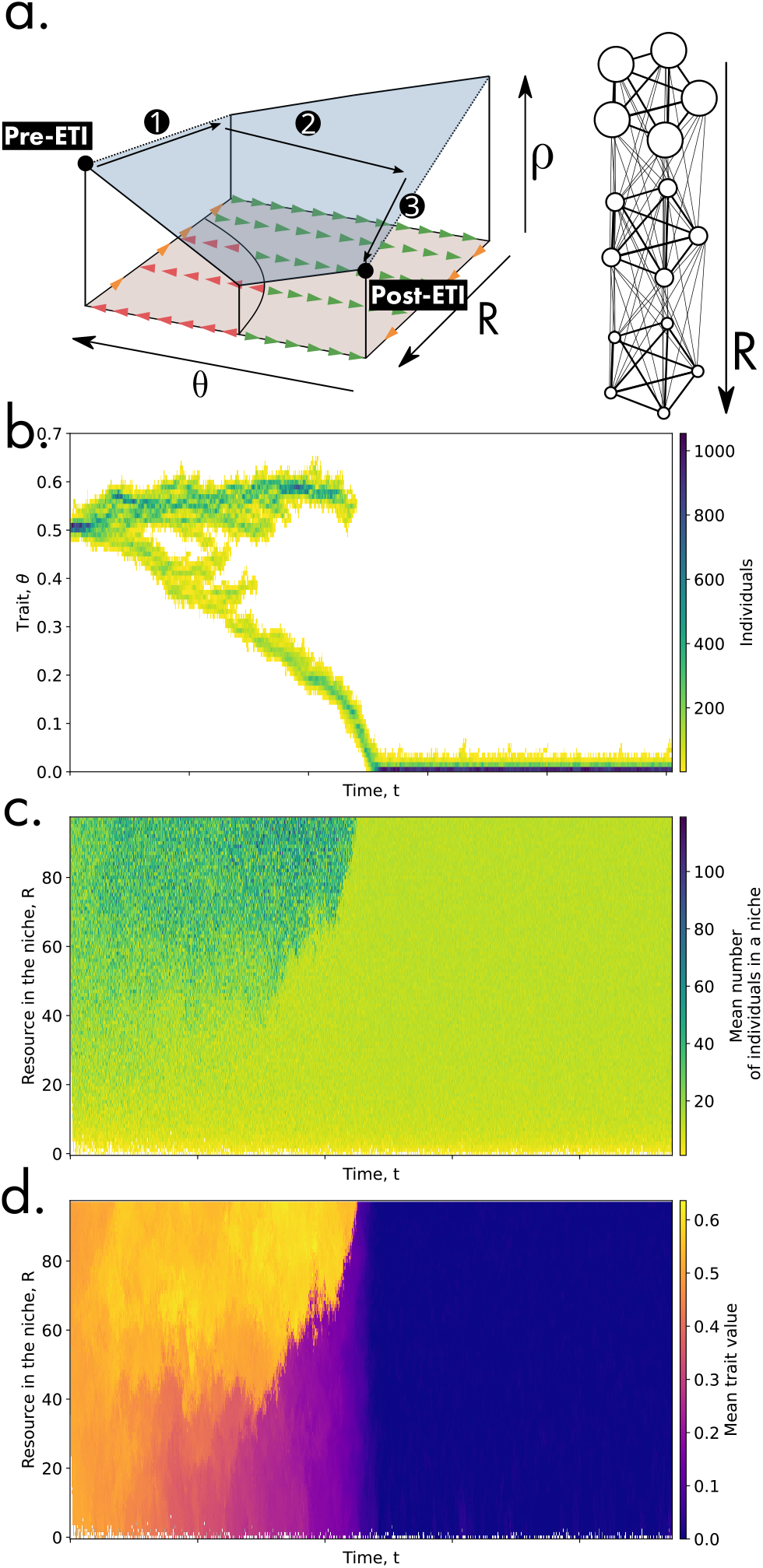
Goldilocks zone. **(a)** *Niche network*. In this new model, the niches are structured along a gradient from resource-rich niches (*R* = 50) to resource-poor niches (*R* = 0). All niches from a single layer are connected to one another as well as to niches of the richer and poorer layers. **(b)** *Trait distribution through time*. The initial trait distribution is centred around *θ* = 0.5. **(c)** *Number of individuals in each category of niches through time*. Initially, only the richest niches (*R* = 50) contain particles. These subsequently migrate down the gradient. **(d)** *Mean trait value in each category of niches through time*. Note how lower values of *θ* first emerge in resource-poor niches and then migrate back to resource-rich niches.

The trajectory can be described as having three crucial phases. First, the population expands to occupy all the niches that are viable (this happens very fast on the evolutionary timescale and is not visible in Figure 5b–d). However, the niches that are too poor in resources (*R < R*_0_) are not populated most of the time because the populations are too small. Second, the populations evolve towards the optimal trait value in each part of the gradient. Two qualitative behaviours exist along the gradient (separated by a fold bi-furcation in *R*^∗^ = 39 as shown in Supplementary Figure 7). For niches where *R > R*^∗^, the populations evolve towards the non-scaffolded peak value *θ*_*u*_ = 0.57, while for *R < R*^∗^, they evolve towards the scaffolded value *θ*_*s*_ = 0. Finally, particles that have evolved the scaffolded phenotype, which is a reliable sensory system, eventually disperse back into the rich niches and out-compete the ancestral type.

Thus, the area in the gradient between the viability limit and the scaffolding limit (*R* such that *R*_0_ *< R < R*^∗^) constitutes a “Goldilocks zone,” where the population structure is suitable for ecological scaffolding. In the children’s tale, only one of the three bears’ belonging is “just right” for Goldilocks, similarly in the diversity of meta-population structures, only some of them are “just right” for scaffolding. As such, the Goldilocks zone acts as a population source (in the ecological source-sink sense) for collectives. The ecological range of collectives (i.e., the portion of the gradient in which their population is stable) is wider than the potentially small Goldilocks zone, which is the only place they can evolve in the first place. On a more general level, it means that the Goldilocks zone can act as an initiator of the ETI.

## Discussion

The notion of (evolutionary) individuality is key to understanding ETIs, as the latter involve a shift in the level at which this property is ascribed. Individuality has been discussed intensively in the philosophical and biological literatures [10, 23–29], with different criteria for characterising what an evolutionary individual is. However, some commonalities can be identified in these different approaches. Discretisation and the evolution of collective traits have been singled out as particularly important aspects of individuality. In this work, we study the case of ecological scaffolding, where these two aspects derive from an externally imposed meta-population structure, and then describe a third aspect: the stability of selected collective traits in the face of changes to the environment (i.e., their endogenisation).

Discretisation is the problem of *group formation* in ETIs [2, 3]. In particular, the population needs to be structured *in discrete collectives* rather than in *neighbourhoods*. Only then can there be genuine collective-level selection as opposed to mere collective-level redescription [30, 31]). Several “individuation mechanisms” for discrete collectives have been proposed [26, 32]. For instance, in nascent multicellularity systems, imperfect cell division and cluster fragmentation give rise to discrete collectives (e.g., snowflakes yeasts, [20, 33], algal clusters [34], cyanobacterial filaments [35]). In ETIs that occur via endosymbiosis, engulfment of one of the particle types provides the template for discrete population structure [15]. In our model, discretisation is established by an externally imposed meta-population structure (following the ecological scaffolding scenario developed in [17, 18]). However, our model allows for overlapping collective generations without compromising simple spatial structure (i.e, by using a graph of niches rather than a continuous resource field). Future work should further explore the parameterisation of the scaffold and create an even more ecologically realistic model. Of particular interest is the effect of mixing between niches, a factor that our model deems negligible. Such mixing could have non-trivial effects on the topology of the fitness landscape and potentially make both scaffolding and endogenisation more difficult.

An individual is also characterised by the evolution of traits that are ascribed at its level. The question of what kind of trait or trait value can be considered truly “collective” rather than a mere aggregation of individual traits has long been the focus of (at times, heated) conceptual discussions (initiated by [36] with the fleet of deer metaphor and followed by [8, 37–39]). A recent approach we follow here tackles this problem by comparing collective-level trait values against linear functions (e.g., averages or sums) of trait values that constituent particles would have if they were solitary. This type of approach can be framed in terms of trait aggregativity versus non-aggregativity [31, 40, 41], counterfactual fitness approaches [42, 43] or indirect genetic effects in quantitative genetics ([44] and [45] chap. 22). Accordingly, two kinds of traits are considered in the present model. In the first case (intrinsic particle dispersal-duplication ratio), the probability of duplicating or migrating is independent of external cues to the particle: a collection of particles only displays an aggregation of identical behaviour. In the second case (density-dependent dispersal-duplication ratio), however, this proba-bility depends on a sensory mechanism that allows synchronisation between cells: their behaviour is not reduced to the average of behaviours as it would be if they were alone. We show that, in scaffolding conditions, the first kind of trait is evolutionarily unstable and is replaced by the second kind of trait. The evolution of synchronisation between different particles through ecological scaffolding represents a step towards greater collective-level individuality as the trait deviates from a purely aggregative one.

While our model features both discretisation and non-aggregative traits at the collective level as necessary features of an ETI, our major result shows that these features of individuality are jointly insufficient for considering collective-level entities full-fledged individuals. We demonstrate not only that collective-level traits can evolve under a scaffold but also that, depending on the conditions under which they evolve, they can remain stable when those conditions change. We take this property of stability to be an important dimension of individuality because a degree of independence between a candidate biological system and its environment features centrally in many accounts of biological individuality. For example, information theory–derived independence from the environment has been proposed to quantify an aspect of individuality [46], while biological autonomy (defined as the ability of organisms to distinguish themselves from their environment) has been recognised as a main concern of evolutionary transitions [47] (see [28] for an exhaustive review). Moreover, the stability of earlier multicellular organisms in the face of single-cell revertants has been identified as a major threat to nascent higher-level entities [7, 48–50]. In stark contrast to the earlier problem of collective formation, this is clearly a problem of collective maintenance [2], associated with mechanisms such as the existence of ratchet mutations [51] or tradeoff-breaking [52]. In the ecological scaffolding scenario, for an ETI to occur, it is not sufficient that collective-level entities are discrete and evolve non-aggregative traits. These non-aggregative traits must also be *endogenised*. If endogenisation does not occur, the collective entities are at best ersatz individuals or, when referring to the traits underlying the capacity for these entities to participate in a selection process, “Darwinian-like” rather than genuinely Darwinian [17, 19, 22]. Our model shows that, given a scaffolding environment, a non-aggregative trait like synchronisation can be endogenised and then invade environments where it was previously unable to evolve.

Our results delineate three necessary conditions for endogenisation: (i) the presence of a fitness valley (ii) that does not exist in the scaffolding environment and (iii) for which the evolutionary endpoint of the scaffolding process must be on the other side of the fitness valley. When these three conditions are met, scaffolding allows collectives to circumvent what would otherwise be an uncrossable fitness valley and then subsequently stabilise the evolved collective-level (i.e., non-aggregative) trait. For illustrative purposes, we have chosen to use a simple picture that combines a one-dimensional environment with a one-dimensional trait value. In principle, however, there is no limit to the number of traits or the number of environmental parameters that could be tracked simultaneously.

These results can help to develop intuitions about experimental systems. Imagine an experiment in which an ancestral population is exposed to candidate-scaffolding conditions for a certain period of time. If no evolutionary response in the collective trait value is observed, new meta-population parameters should be tested. If a reversal of the collective trait is observed upon return to the ancestral environment, this could be because the duration of scaffolding was not sufficiently long to enable crossing of the fitness valley (this could potentially be resolved by maintaining the scaffolding regime for longer), or possibly because there was no fitness valley at all (this could be tested by looking for qualitative changes in the evolutionary response when trying many different meta-population parameters; see Supplementary Figure 6).

The ecological scaffolding model has been successfully used to interpret evolutionary experiments where the environment is fully controlled by the experimenter [11]. Understanding what it can and cannot do, as well as the effect(s) of meta-population structure parameters, is of empirical interest for the fundamental study of ETIs and for applications in the field of artificial selection of communities [18, 53–56]. In particular, the stability of selected traits beyond the environments that promoted their initial emergence is a highly desirable property for engineered collectives [57]. Three increasingly strong notions of trait stability might be of interest in this context. First, a trait might be ecologically unstable, disappearing in one or few generations or being replaced by existing variants in the population. Alternatively, a trait can be evolutionarily unstable (as in [18]), meaning that the trait would disappear in the long run due to the successive invasion of new mutants. Last, a trait might be evolutionarily stable (e.g., as a result of endogenisation), meaning that small mutations would not eventuate in replacement. The last case is the most desirable in applied settings because invading phenotypes that do not contribute to the trait of interest pose a serious threat to synthetic consortia.

A key outcome of our work is that not all externally imposed meta-population structures lead to the scaffolding and endogenisation of collective-level traits. A direct implication is the possibility of zones that can be considered causally operative factors or “ETI initiators,” which, for hopefully obvious reasons, we have chosen to call “Goldilocks zones.” Indeed, it is possible that along an environmental gradient, only some areas meet the conditions for the processes of scaffolding and endogenisation (in the Goldilocks tale, this would correspond to the one bear’s belongings that enable “just the right” set of affordances for their trespasser’s prolonged stay). When this happens, pre-ETI particles can immigrate into the Goldilocks zone, be selected for collective organisation, and then emigrate out of the zone into another environment. For example, it has been hypothesised that the origin of the first cells relied on the iron-sulfide crystalline structure in hydrothermal vents [58–60]. That particular structure could act as a Goldilocks zone and thus have featured as the source of the cells that have since colonised our entire planet. Endogeni-sation of traits would be required for this to happen. An additional condition of the gradient-scenario compared to the limited-duration scaffolding scenario described above is the non-competitive exclusion of the derived type by the ancestral type (so the new collectives can invade back the ancestral environment) or that the newly formed collectives colonise parts of the environment where ancestral particles were not viable. This scenario furnishes us with an especially intriguing consequence: Goldilocks zones can apparently be relatively rare in the overall environment without thereby foreclosing or even limiting the possibility of ETIs, not unlike hydrothermal vents that occupy only a small fraction of the ocean’s floor. The ecological range of scaffolded collectives that are subsequently endogenised can be broader than the (possibly limited) range of environments that promote their emergence in the first place.

## ACKNOWLEDGEMENTS

The authors gratefully acknowledges the financial support of the John Templeton Foundation (#62220). The opinions expressed in this paper are those of the authors and not those of the John Templeton Foundation.

## Author Contribution

All authors conceived the study. GD wrote the model and performed the analysis. All authors contributed to writing the manuscript. KH and PB contributed equally.

## Supplementary Note 1: Eco-evolutionary Meanfield Niche Model

In this section we study the meanfield model approximating the eco-evolutionary dynamics of the meta-population. We establish the invasion fitness gradient.

### A. Description

Let *C*(*t*) be the density of occupied niches at time *t* and *E*(*t*) be the density of empty niches at time *t*. Consider that *C* and *E* follow the “SIS” dynamics:

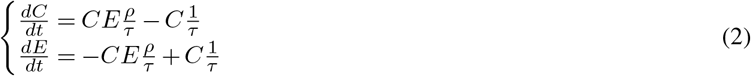

Initial conditions will be (1, *D*), with *D >* 0.

### B. Ecological equilibrium

The ecological equilibrium is reached when 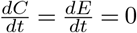. There are two branches of equilibria:

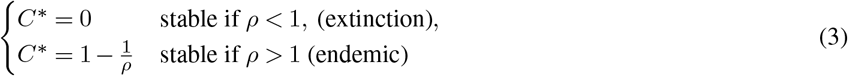

**Sup. Fig. 1.**
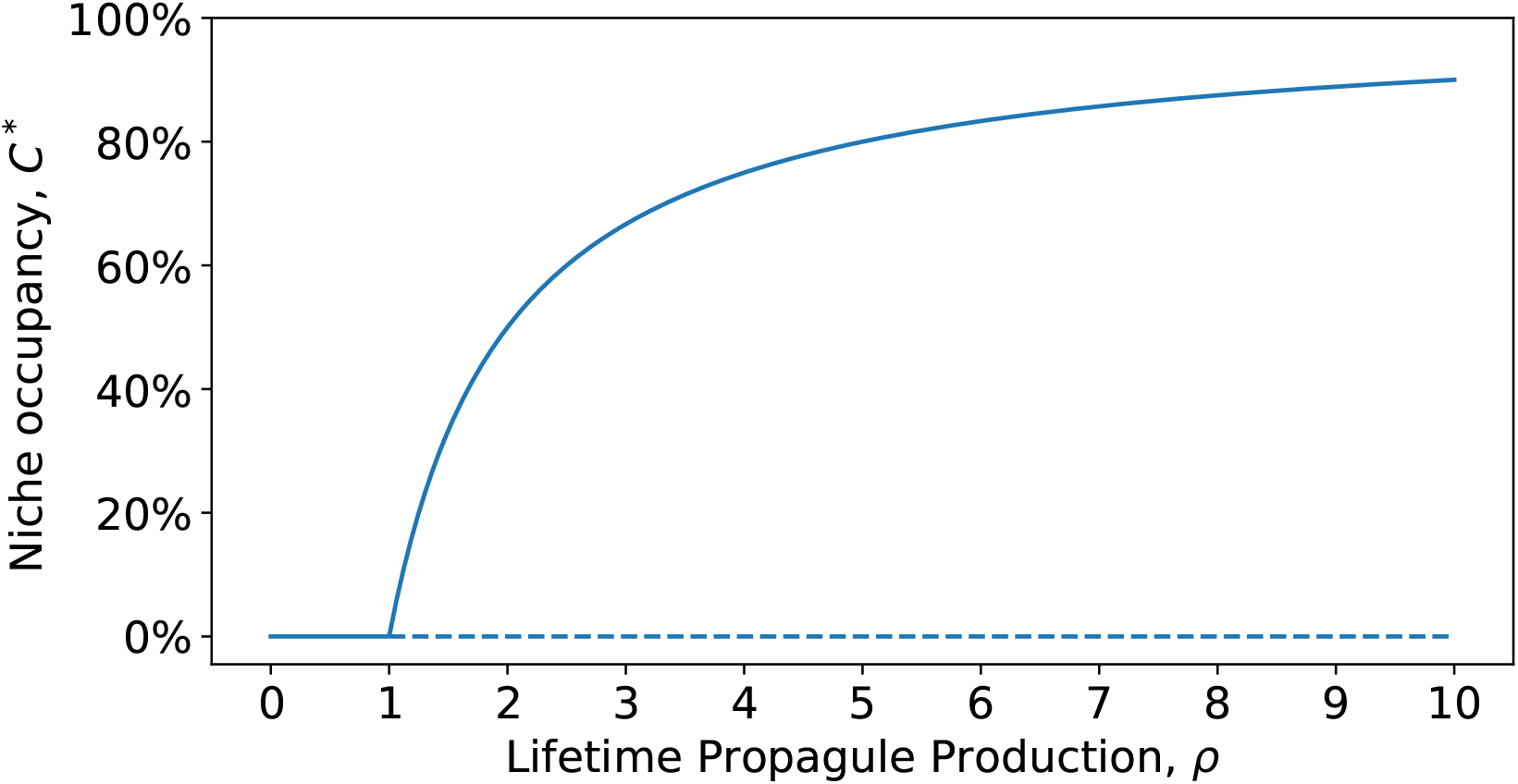
Niche occupancy at equilibrium *E*^∗^ as a function of the average propagule count.

In both cases, *E*^∗^ = *D* − *C*^∗^. Note that there is a transcritical bifurcation in *ρ* = 1 (Supplementary Figure 1).

### C. Adaptive dynamics

Let 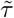 and 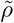 be functions of an underlying trait *θ*, such that 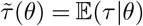 and 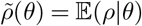. First, we compute the invasion fitness from the ODE model by adding a mutant with a trait *m* in a population with resident trait *r*:

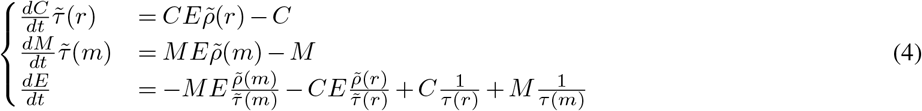

And looking for the exponential growth rate of a rare mutant in a resident population at non-extinct equilibrium—that is, 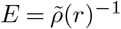 and 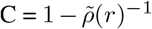:

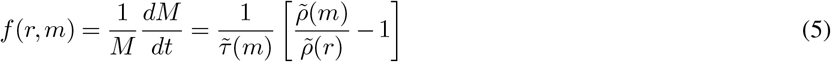

Supplementary Figure 2 shows the sign of the invasion fitness *f* as a function of *θ* for three values of *R*, taking the simplest functional form for *p* as a function of the trait *θ*, taking *p*(*θ*) = *θ* (referred to as the intrinsic constant duplication-dispersal ratio in the main text).

The invasion fitness gradient *g* (i.e., the derivative of the invasion fitness evaluated when the mutant trait is equal to the resident trait) is:

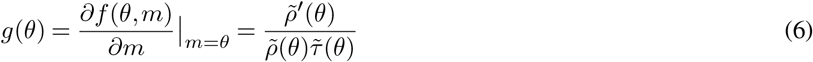

**Sup. Fig. 2.**
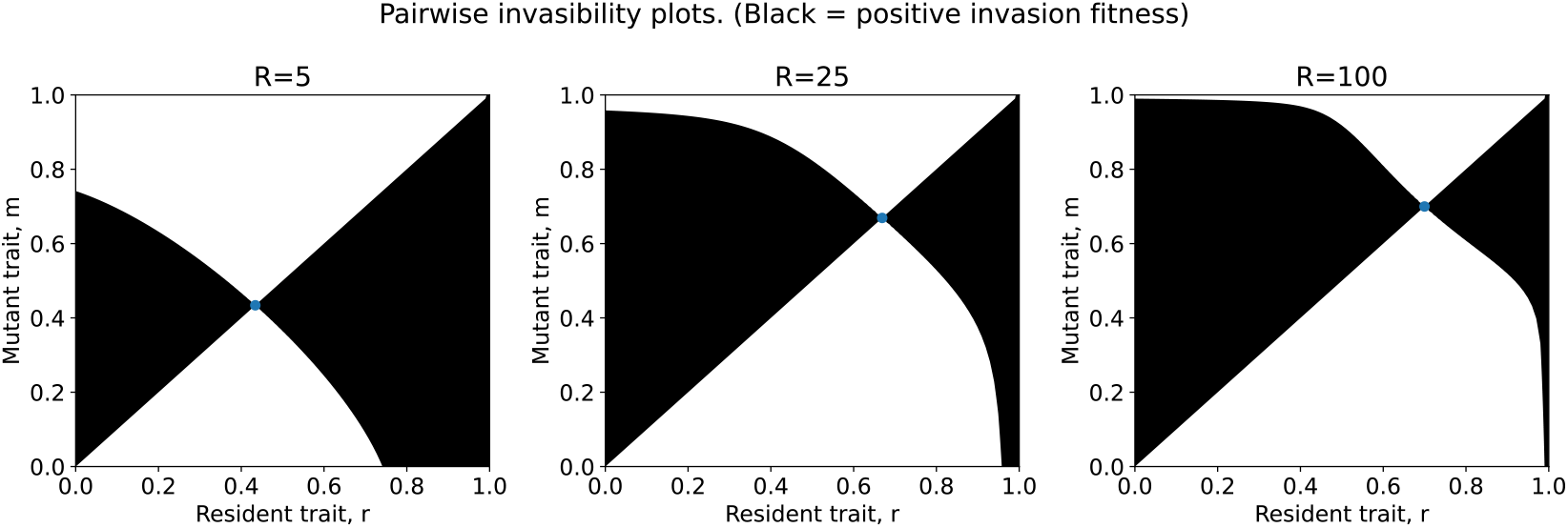
Pairwise invasibility plots for the intrinsic constant duplication-dispersal ratio. In black are values of resident *θ* = *r* and mutant trait *θ* = *m* that correspond to a positive invasion fitness *f* (*r, m*) and thus invasion by the mutant; in white are values that correspond to a negative invasion fitness and thus no invasion by the mutant. Invasions by a mutant with an infinitesimal trait difference are studied by looking at the sign changes along the first diagonal. A singular value (blue dot) occurs where the sign changes. In the three panels presented, it is always convergent and evolutionarily stable (See [61] for more details about the method of Pairwise Invasibility Plots).

In the approximation we are making (i.e., rare mutant, resident at ecological equilibrium), the lifespan *τ* does not matter for the endpoint of the evolutionary trajectory outside the lifetime reproductive output *ρ*. The ESS is simply the trait value that optimises *ρ*. This can be seen in the expression of *g*. Since *τ* and *ρ* are always strictly positive, the values of *θ* for which *g* is null are the values of *θ* for which *ρ*^*′*^ is null, and hence the values of *θ* for which *ρ* is maximal (or minimal).

Supplementary Figure 3 shows the singular trait value in the intrinsic constant duplication-dispersal ratio model obtained by numerically computing the root of the function *g*(*θ*) or, equivalently, the maximum of the function *ρ*(*θ*). (For the ESS in the density-dependent duplication-dispersal model, see Supplementary Figure 7).

**Sup. Fig. 3.**
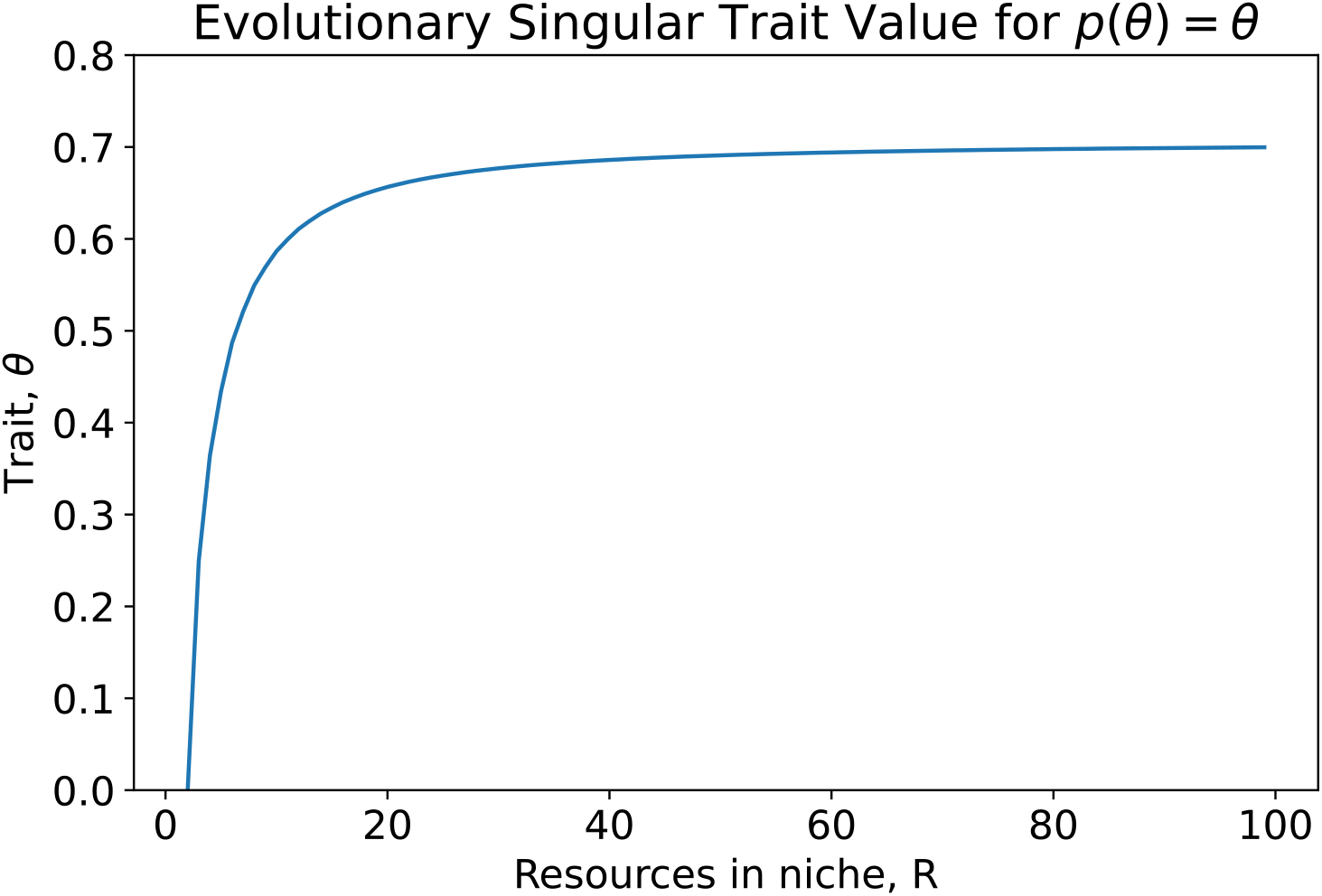
Evolutionarily singular strategies for the intrinsic constant duplication-dispersal ratio.

One simplification of the modelling work we present is rooted in the adaptive dynamics approach, which supposes a separation of timescale between the introduction of new mutations and their fixation in the population. This limits the ability of this kind of model to quantitatively predict the duration *T* that would be required for scaffolding experiments to be successful. However, provided the fitness landscape has the same topology, qualitative results should hold.

## Supplementary Note 2: Stochastic Niche Model

In this section, we study the stochastic model of particle dynamics within a niche and derive the expression for the expected life span and the expected number of propagules.

### A. Constant probability p

Let (*X*_*t*_, *t* ≥ 0) be a one-dimensional continuous time Markov branching process modelling the number of particles in a niche. Let *R* be the number of resources in a niche, *υ* be a timescale factor (set to 1 in the main text for easier presentation), and *p* be the probability to stay in the niche on a splitting event or send out a single propagule.

(*X*_*t*_, *t* ≥ 0) is a linear birth-death process with per capita birth rate *υp* and per capita death rate *v*(1 − *p*). The process is stopped once all particles are dead (*X*_*t*_ = 0) or at the *R*-th splitting event (i.e., when the resources are depleted).

We are looking for the lifespan of the collective *τ* and the number of propagules *ρ*.

#### A.1. Expected values

The branching process (*X*_*t*_, *t* ≥ 0) can be separated into two sub-processes: a jump process (*S*_*n*_, *n* = 1 … *R*) and a split time process (*T*_*n*_, *n* = 1 … *R*) [62, p. 118]. *S*_*n*_ is the number of cells at the n-th split time, and *T*_*n*_ is the time of the n-th split.

##### Jump process

The jump process (*S*_*n*_, *n* = 1, 2…*R*) is a discrete time random walk, starting at *S*_1_ = 1 and stopped if it reaches 0, with transition probabilities for *i >* 0 given by:

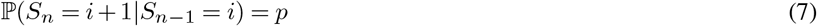

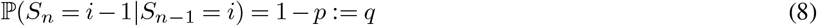

Consider a jump *n* such that 1 *< n < R*—let us compute the discrete probability distribution *P* (*S*_*n*_ = *k*) for all values of 0 *< k < R*.

**First case**, *k* ≥ 1 A trajectory of *S* is a path from (*t, x*) = (1, 1) to (*k, n*) that does not cross the axis (*·*, 0).

- It can be proven by reflection that there are 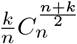 of such paths (the Ballot theorem).
- A path from (1, 1) to (*k, n*) is composed of (*n* + *k*)*/*2 − 1 steps up and (*n* − *k*)*/*2 steps down. The probability of any such path is 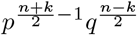.

The probability ℙ (*S*_*n*_ = *k*) is the product of this probability of getting any path from (1, 1) to (*k, n*), with the number of such path. Thus:

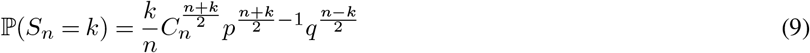

**Second case**, *k* = 0

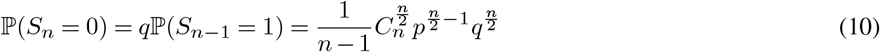

##### Number of propagules

There are as many propagules produced for a given trajectory of *S* as there are steps down in the corresponding path. Thus:

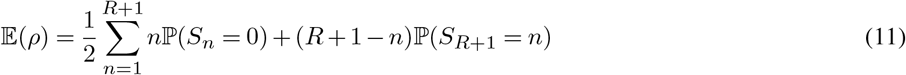

##### Split time process

Conditionally to the trajectory *S*, the split time process is a series of independent exponential random variables:

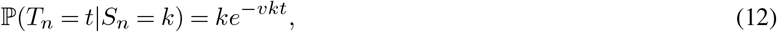

for *k >* 0. In the case *k* = 0, *T* is deterministic and equal to 0.

This means that the conditional expected value is 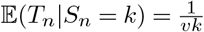.

The life span of a collective *τ* is given by the sum of all the splitting times *T*. Thus:

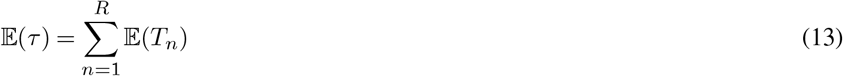

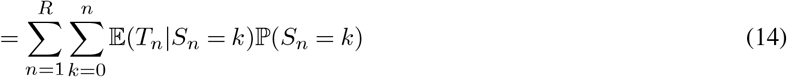

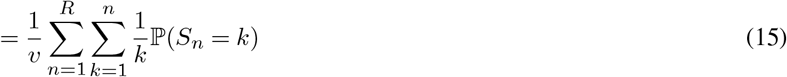

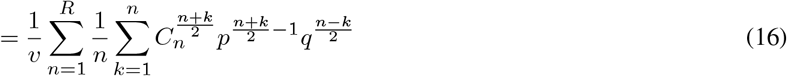

#### A.2. Special values

For *p* = 0, the first event is always a particle dispersal:

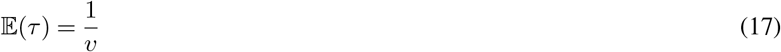

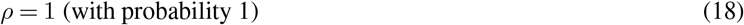

For *p* = 1, no propagule is produced and the collective lifespan is just the harmonic series:

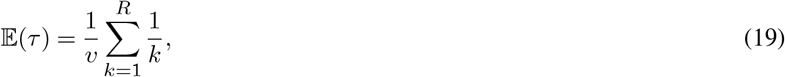

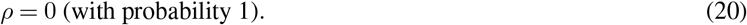

### B. Density-dependant *p*

Let *p*: *n* → *p*(*n*) be a function of *n*, the number of particles in the niche.

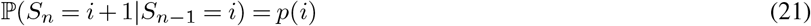

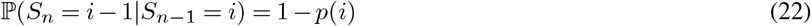

Let *M* be the transition matrix of the process *S*, such that 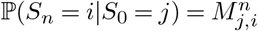. From Equation 11, we have:

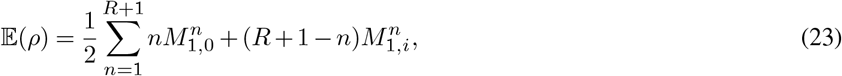

with

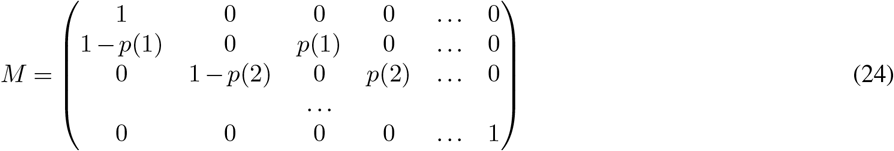

## Supplementary Note 3: Stochastic simulations

In this section, we describe the algorithm used for the stochastic simulations of the meta-population eco-evolutionary dynamics. A simulation of the model takes as input:

- A *scaffold*, composed of an undirected graph *S* = (*V, ℰ*) where each vertex is a niche, and a function *ℛ* that associates each niche to a number of resources. An edge between two vertices means that propagules from one niche can reach the other niche.
- A *trait space T* that encodes the possible trait values, and a mutation rate *μ*. Particles inherit the trait value of their parent with probability (1 − *μ*) or mutate to an adjacent value in the trait space with probability *μ*.
- A *particle ecology* function that associates the trait value of a focal particle *θ* ∈ *T*, the number of available resources *R* ∈ ℕ, and the number of other particles in the niche *N* ∈ ℕ to the probability that a particle would duplicate or form a propagule. *p*: *θ, R, N* ↦ *p*(*θ, R, N*)
- An *initial population* of particles: that is, a number of particles with associated trait value and niche location.
- A *duration* of the simulation *T* ∈ ℝ_+_.

We use the exact stochastic simulation algorithm (SSA) to generate the trajectory of the system, starting from time *t* = 0, and while *t < T* the time to the next event is sampled from an exponential distribution, and the nature of the next event is sampled from a multinomial distribution with appropriate parameters. The possible events are, for each particle:

- Particle birth, with rate *p* for each particle. Add a particle to the same niche, reduce the number of resources in the niche by 1. With probability *μ*, the new particle mutates to a different trait value than its parent.
- Propagule production, with rate 1 − *p*. Remove the particle, reduce the number of resources in the niche by 1. Select uniformly at random a niche adjacent to the current niche. If this niche does not contain any particle, add a new particle with the same trait value to this niche.

If after applying the event, the number of resources in the focal niche *e* is 0, reset the niche: remove all particles and replenish the resources to its initial value *ℛ*(*e*).

- In the intrinsic particle dispersal-duplication ratio, the ecology function is simply the constant function *p*: *θ, R, N* ↦ *θ*.
- In the density-dependent particle dispersal-duplication ratio, the ecology function is:

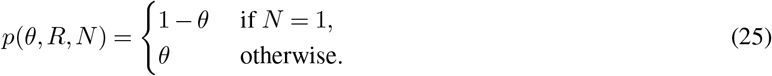

## Supplementary Note 4: Evolutionary trajectories

In this section, we check that the meanfield approximation is coherent with the stochastic simulations. We start with a complete network (i.e., each niche is connected to each niche), and constant resource richness. Supplementary Figure 4 shows stochastic ecological trajectories. Note that the stochastic occupancy corresponds to the prediction of the meanfield model.

**Sup. Fig. 4.**
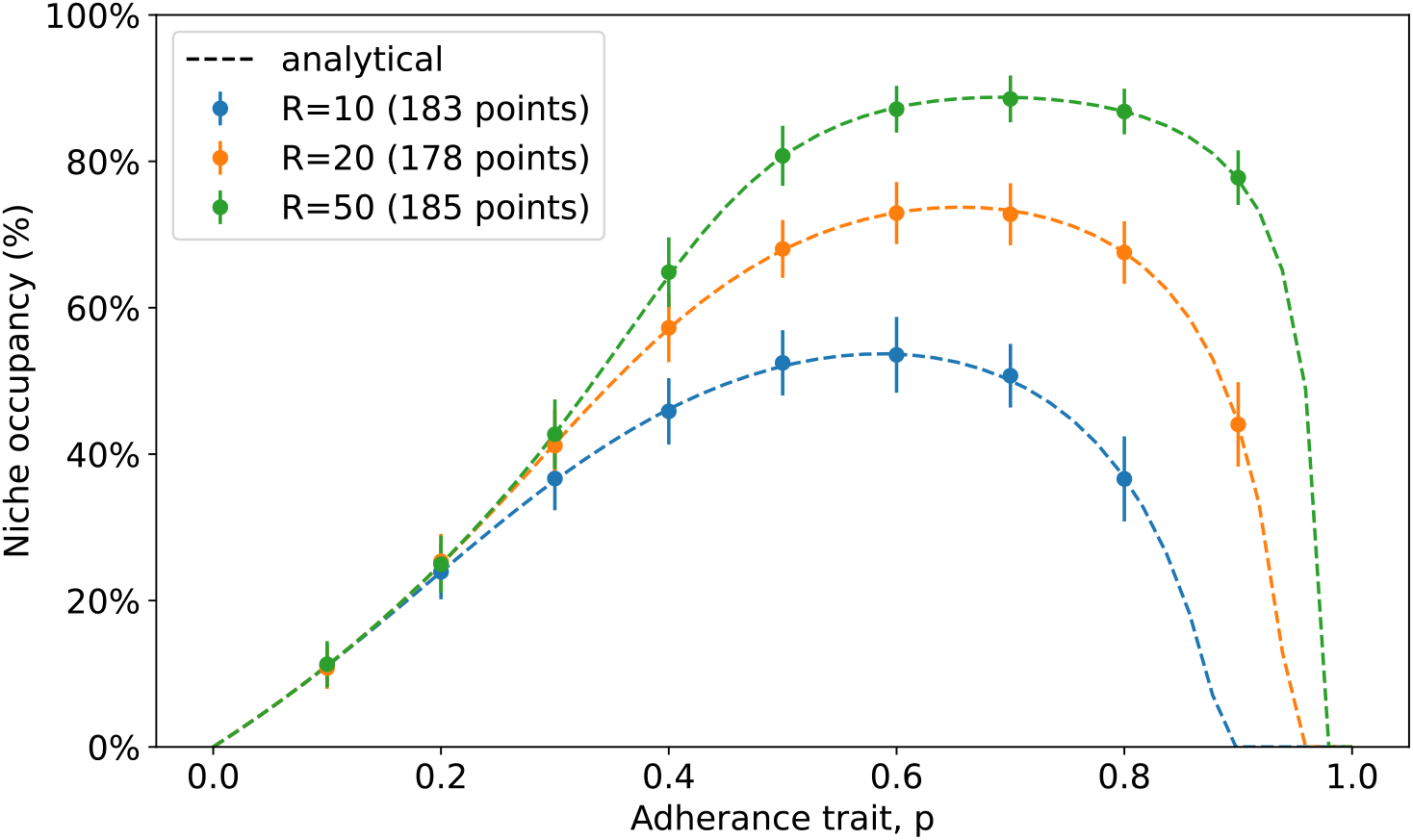
Occupancy with constant scaffolding conditions. Proportion of niches that are occupied as a function of *θ* in the intrinsic duplication-dispersal ratio. The scaffold is a complete graph of *D* = 100 niches with a uniform number *R* of resources, and no mutations are allowed. The line is the prediction of the meanfield model. Each dot is the average occupancy in stochastic simulation (number of replicates in the legend).

Supplementary Figure 5 shows the life span and number of propagules sent during the life span of the niche. Note that the expected value computed in Supplementary section 2 is a good approximation of the results of the stochastic trajectories.

Supplementary Figure 6 shows the evolutionary trajectory obtained by stochastic simulations. Note that the endpoint of the simulations corresponds to the prediction of the meanfield model.

**Sup. Fig. 5.**
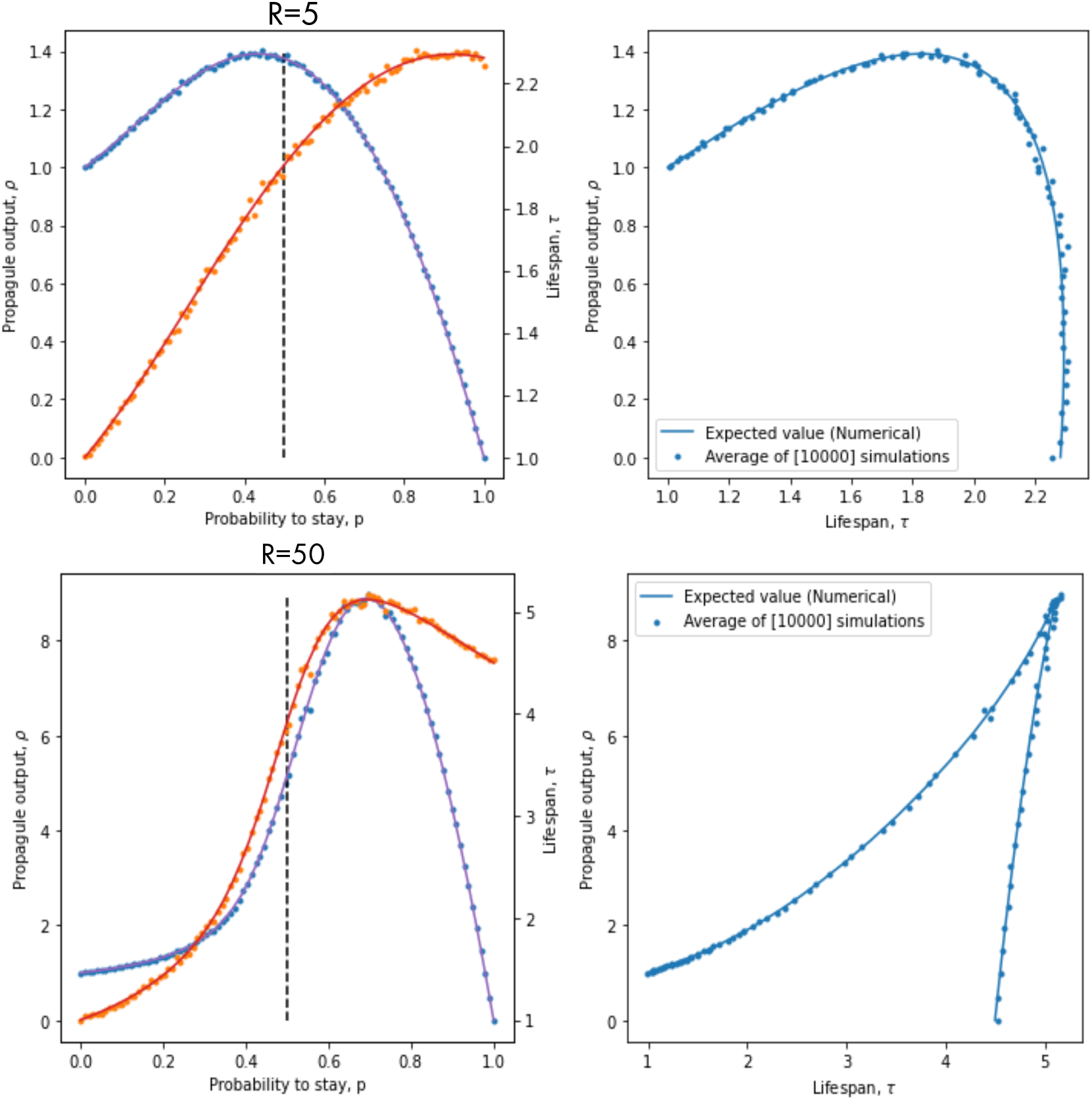
Single collective simulations. Only a single collective is simulated, with a number *R* of resources and a trait value *θ* in [0, 1] for the intrinsic ratio.

## Supplementary Note 5: Conditions for endogenisation

Consider two environmental values, called scaffolding *e*_*s*_ and non-scaffolding *e*_*u*_. Now, 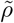 and 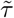 are functions of the underlying trait *θ* and the environment *e*. We assume that they are continuous.

Endogenisation relies on the existence of a hysteresis cycle as follows, when the four following conditions are fulfilled.

- **Condition 1 (Non-scaffolded ESS)**. There exists a pre-ETI trait state *θ*_*u*_ and a post-ETI state *θ*_*m*_, which are both evolutionarily stable in the non-scaffolding environment (the subscript stands for “unicell” and “multicell”). Without loss of generality, assume that *θ*_*u*_ *< θ*_*m*_. It means that the function 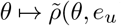) has two local maxima in *θ*_*u*_ and *θ*_*m*_. Since 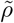 is continuous, there exists a local minimum *θ*_−_ such that *θ*_*u*_ *< θ*_−_ *< θ*_*m*_. Namely:

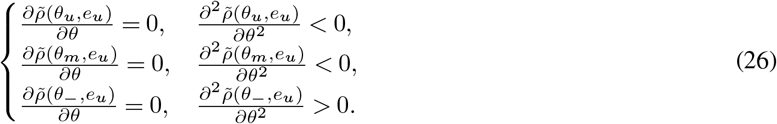
- **Condition 2 (Scaffolded ESS)**. There exists an evolutionarily stable state *θ*_*s*_ in the scaffolding environment. Namely:

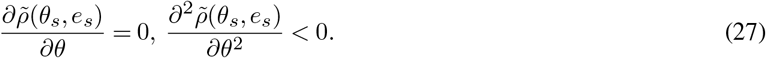 For simplicity, we will consider that it is unique, but it would be enough to have its basin of attraction include both *θ*_*u*_ and *θ*_*m*_.
- **Condition 3 (Ecological stability)**. The population must be viable for the 3 ESS, namely:

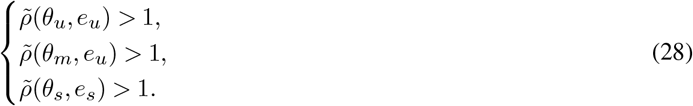
- **Condition 4 (ESS order)**. The scaffolded state must be in the basin of attraction of the post-ETI ESS. Namely:

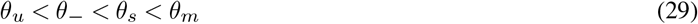

Supplementary Figure 7) illustrates how the density-dependent duplication-dispersal ratio model fulfils these conditions.

**Sup. Fig. 6.**
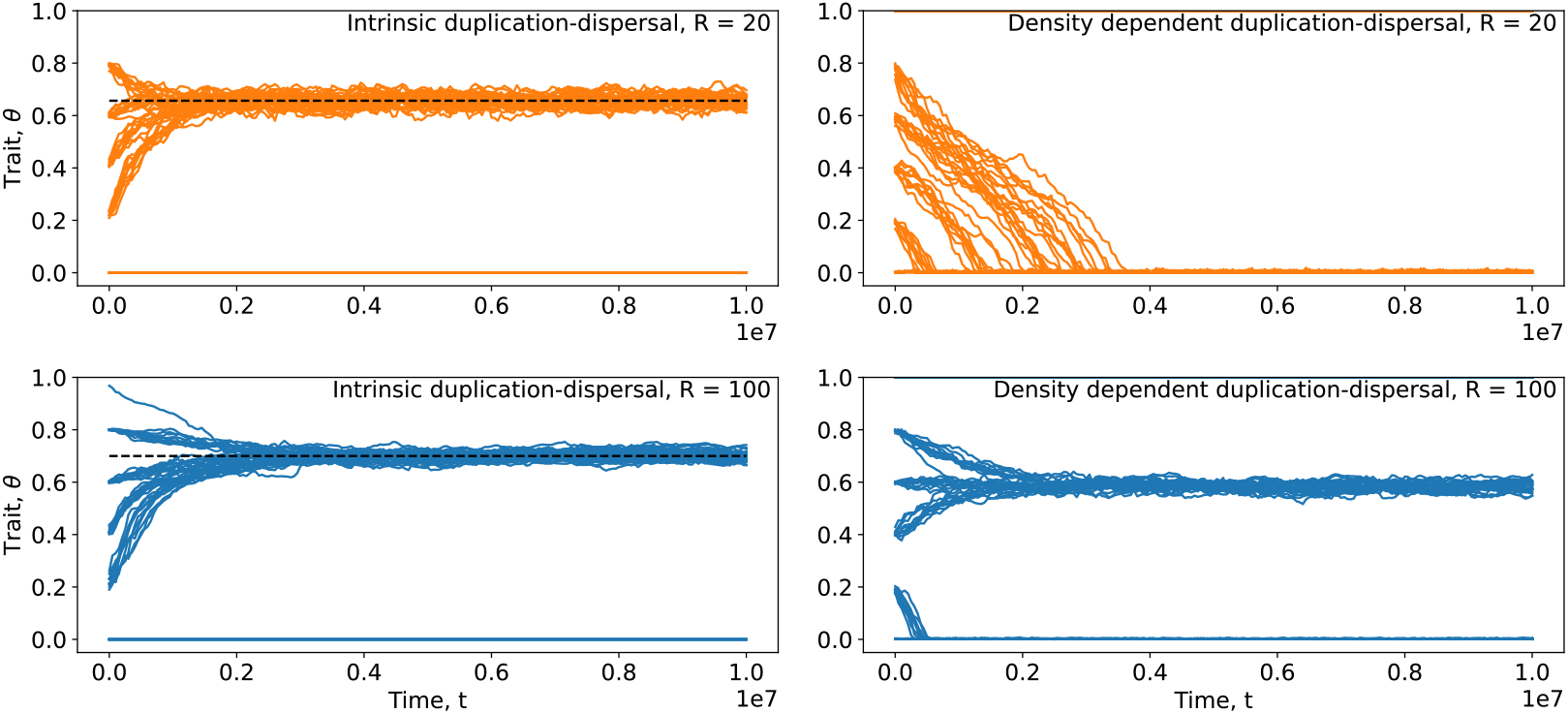
Evolutionary strategies with constant scaffolding conditions. In these simulations, the scaffold is a complete graph of *D* = 100 niches with a uniform number *R* of resources and an initial trait value *θ*_0_ in (0, 0.2, …0.8, 1) for both ratio functional forms. Each line is an independent stochastic simulation, and 10 replicates are simulated for each set of conditions. The dotted line corresponds to the adaptive dynamics prediction in the meanfield model.

**Sup. Fig. 7.**
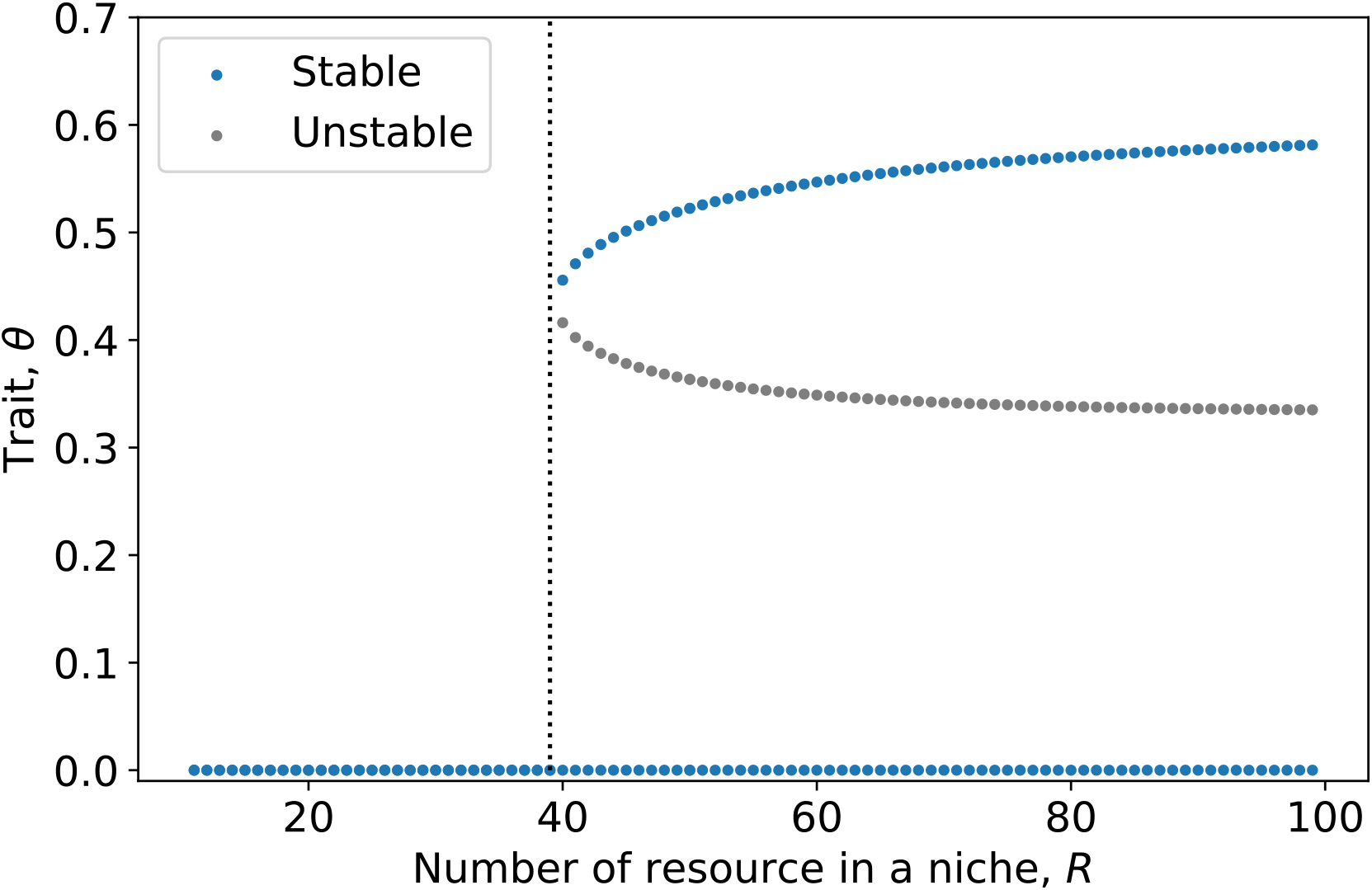
Singular Evolutionary strategies for the density-dependent duplication-dispersal ratio as a function of *R*. There are three branches of equilibria. Note the fold bifurcation at *R* = 39, the stable branch corresponding to an imperfect sensory system (*θ*_*u*_, top-right, stable) collides and annihilate each other with the bottom of the fitness valley (*θ*_−_, middle, unstable). The perfect sensory system (*θ*_*m*_ = 0, bottom) is always stable. If the system is on the top branch (*θ*_*u*_ *>* 0.5, *R* = 50), a change from *e*_*u*_ : *R* = 50 to *e*_*s*_ : *R* = 20 causes the system to irremediably move to the bottom branch (*θ*_*m*_, coordination between cells). If the environment is restored to *R* = 50, the population does not revert to the ancestral phenotype. Such an irremediable change is called hysteresis in ecology [63].

